# Parity modulates impact of BMI and Gestational Weight Gain on gut microbiota in human pregnancy

**DOI:** 10.1101/2022.09.02.506244

**Authors:** Katherine M. Kennedy, Andreas Plagemann, Julia Sommer, Marie Hofmann, Wolfgang Henrich, Jon F.R. Barrett, Michael G. Surette, Stephanie Atkinson, Thorsten Braun, Deborah M. Sloboda

## Abstract

Pregnancy requires maternal adaptations to support fetal growth: whether these adaptations include temporal shifts in the gut microbiome is still unclear. We investigated the maternal gut microbiome longitudinally over the course of pregnancy and the impact of pre-pregnancy BMI (pBMI) and GWG. We also determined whether parity modulated observed associations. We show that the gut microbiota of participants with higher pBMI changed less over the course of pregnancy in primiparous, but not multiparous participants. This suggests that previous pregnancies may have persistent impacts on maternal adaptations to pregnancy. This ecological memory appears to be passed to the next generation, as parity modulated the impact of maternal GWG on the infant gut microbiome. This work supports a role for the gut microbiome in maternal adaptations to pregnancy and highlights the need for longitudinal sampling and accounting for parity as key considerations for studies of the microbiome in pregnancy and infants.

## Background

Pregnancy requires physiological, metabolic, and immune adaptations to support fetal growth and development^1^. Pregnancy-induced metabolic adaptations mirror those seen in non-pregnant metabolic syndrome; including hyperglycemia, hyperinsulinemia, and increased adiposity. When combined with high pre-pregnancy body mass index (pBMI) or excess gestational weight gain (GWG) these adaptations can often reach unhealthy levels. Indeed, high pBMI is a known risk factor for maternal and infant health complications^2^. Although some data suggest that pregnancy-associated metabolic adaptations may influence^3^ and/or be influenced by^4^ shifts in the gut microbiota over the course of pregnancy, most studies have investigated only one (sometimes two) timepoints during gestation^4–7^. An evaluation of whether gut microbiota change temporally across pregnancy, birth and the postnatal period is required to robustly assess whether host-microbe interactions could govern pregnancy adaptations. Of note, we currently lack even a foundational understanding of whether the maternal microbiome does indeed change over the course of pregnancy.

One of the first studies of the gut microbiome during pregnancy reported an increase in diversity between samples (beta-diversity) and a decrease in diversity within samples (alpha-diversity) in healthy pregnant individuals during the third trimester of pregnancy compared to the first^4^. In contrast, others have found the maternal gut microbiota to be stable over the course of pregnancy^8–10^. Few studies have investigated microbial shifts in the context of pregnancy complications like high pBMI or excess GWG^11–13^. Although excess GWG is more likely to occur in cases where maternal pBMI is high, 50% of all pregnant individuals experience excess GWG. Thus, these two factors both independently and together influence maternal and infant health. To date, no human studies have investigated whether pregnancy-associated microbiotal shifts are influenced by previous pregnancies (parity).

Both high maternal pBMI and excess GWG are associated with pregnancy complications^2^ and impaired offspring metabolism^14,15^. Reports suggest that some of these maternal and infant associations may be mediated by obesity-associated “dysbiosis” of the maternal gut microbiome during pregnancy^7,11^. As humans are not colonized by microbes before birth^16^, the maternal microbiome may influence offspring development via changes to the *in utero* environment (e.g. altered maternal metabolic adaptations to pregnancy) and/or altered seeding of the infant microbiome during and after birth. In this prospective, observational cohort study we set out to determine whether temporal changes in maternal gut microbial composition occur with advancing gestation and the postpartum period. We also investigated whether maternal BMI and/or excess GWG and parity impact maternal microbiotal adaptations and how these contexts influence the maternal and infant microbiomes at 6 months of age.

## Results

### Study cohort

A total of 89 participants were recruited; after we excluded participants who reported any nicotine or antibiotic use during pregnancy, 65 participants were included in the study (primiparous n=40; multiparous n=25). The majority of participants with a pre-pregnancy BMI (pBMI) of 25-30 or >30 had gestational weight gain (GWG) above the Institute of Medicine (IOM) recommended range^17^ (83% and 86% respectively), compared to less than half of participants with a pBMI 18.5-25 (33%; Extended Data Table 1). Most participants delivered vaginally (79%), and mode of delivery was similar across pBMI categories (q=0.8). Multiparous participants had either one (88%) or two previous pregnancies (12%). Multiparous participants were older than primiparous participants (32.8±3.1 vs 29.6±4.4, q=0.009), but there was no significant effect of parity or maternal age on pBMI, GWG, post-partum weight retention, length of gestation, or delivery mode.

### Parity modulates the impact of pBMI and GWG on maternal gut microbiota

Maternal stool samples were collected longitudinally: in the first (10-17 weeks gestation), second (26-28 weeks gestation), and third trimesters (>36 weeks gestation) of pregnancy, at delivery, and at 6 months postpartum. We assessed the maternal gut microbiota using 16S ribosomal RNA gene sequencing of the combined V3-V4 region. We identified minimal sporadic contamination in negative controls, likely originating from samples. Of 13 sequenced gDNA extraction negative controls, 4 had no reads and an additional 4 had fewer than 20 reads. The 5 remaining negative controls had 45, 55, 80, 130, and 384 reads: 65 of 67 ASVs were detected in a single negative control (*Escherichia/Shigella* and *Bacteroides* were each detected in 2 negative controls). These negative control data show no consistent contamination signal. We identified a total of 7577 ASVs in maternal samples after removal of host-associated reads (Kingdom Eukaryota, Order Chloroplast, Family Mitochondria, or no assigned Phylum) and the median sample read count was 62,853 (min = 14,872; max = 136,000).

Alpha diversity was positively correlated with pBMI (R^2^ = 0.11, p=0.010), particularly in participants with GWG above the recommended range overall (R^2^ = 0.12, p=0.0069) and at each pregnancy time point (R^2^ = 0.14-0.21, p=0.012-0.046; Extended Data Fig. 1), but not at 6 months post-partum (R^2^=0.014, p=0.57). To assess differences in overall community composition between samples (beta-diversity) we performed principal coordinate analysis (PCoA) of Bray-Curtis dissimilarity (Fig. 1a, b). Beta-diversity was modestly impacted by GWG category (R^2^ = 2.6%, p=0.0001; Fig. 1a), pBMI category (R^2^ = 2.8%, p=0.0001; Fig. 1b), parity (R^2^ = 1.4%, p=0.0001; Extended Data Fig. 2a), and sample time point (R^2^ = 1.0%, p=0.0001; Extended Data Fig. 2b). These modest effects were overshadowed by the impact of interindividual variability, which explained the majority of variation between samples (PERMANOVA; R^2^=62.3%, p=0.0001). Longitudinal sampling within individuals is necessary to disentangle these large effects of inter-individual variability^18^ from factors impacting pregnancy-associated shifts in the microbiota (intra-individual variability). We assessed beta-diversity longitudinally across time points (Bray-Curtis dissimilarity) *within* individuals and found that the impact of pBMI differed by parity. Beta-diversity was negatively correlated with pBMI at every time point in primiparous (R^2^ = 0.041-0.19, p < 0.0001-0.029) but not multiparous participants (R^2^ < 0.0001-0.016, p = 0.34-0.99; Fig. 1c). These data suggest that in primiparous (first) pregnancy, the gut microbiota is more stable in individuals with a high pBMI over the course of pregnancy and postpartum. Together these observations illustrate that when considering temporal changes in microbial composition related to pregnancy, longitudinal sampling within individuals is essential in uncovering patterns of change that exist with advancing gestation and persist postpartum. These data also highlight a novel observation; that gut microbial diversity during pregnancy and postpartum is influenced by previous pregnancies. Parity therefore is a key confounder that must be considered in all future studies investigating the maternal microbiome.

**Figure 1.**
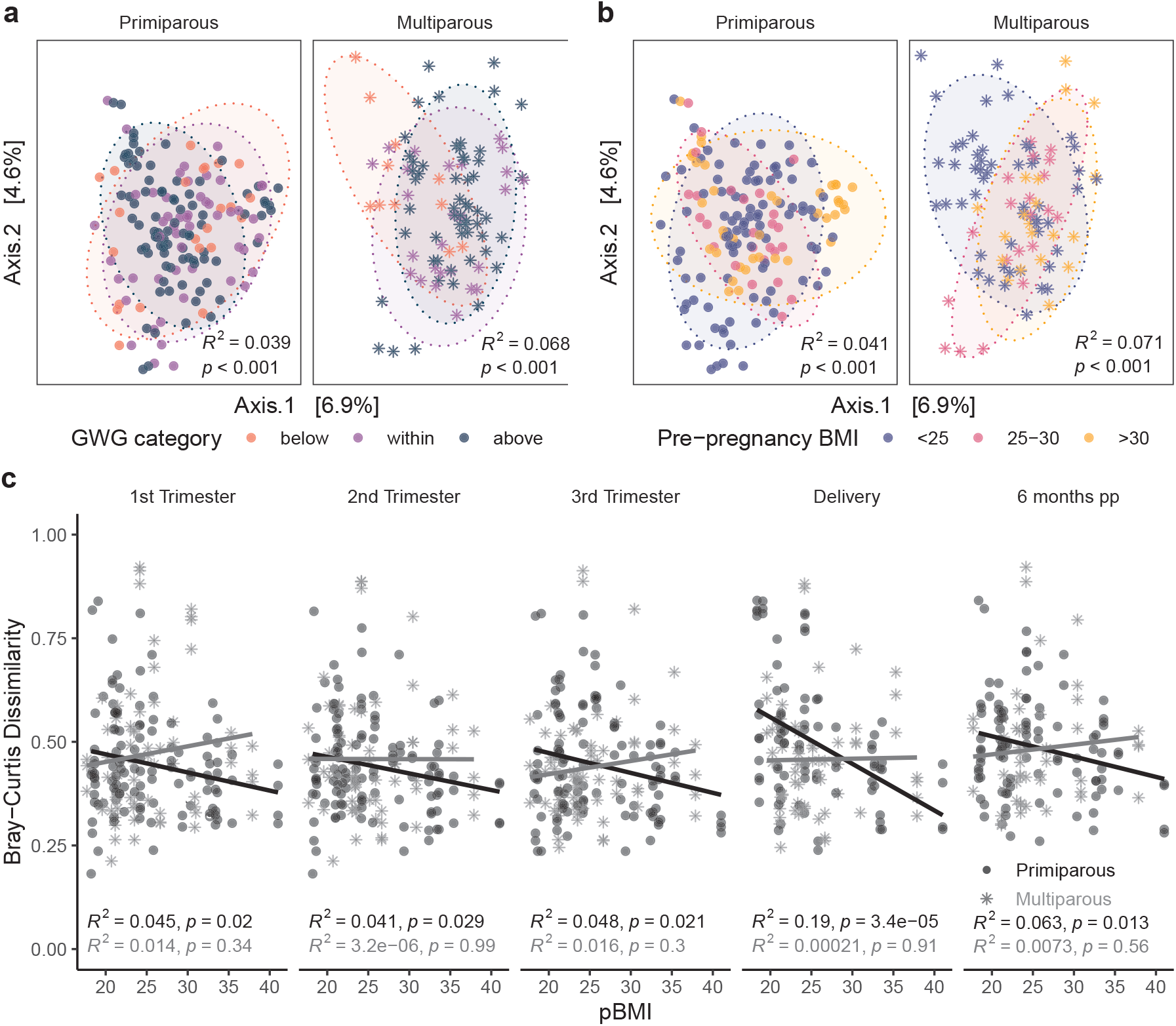
Parity modulates impact of pBMI and GWG on pregnant gut microbiota. PCoA of Bray-Curtis distances in primiparous (n=39, one excluded - no GWG data) and multiparous (n=23, two excluded – no GWG data) participants shows modest clustering by **a**, GWG category (primiparous, R2=0.039, p<0.0001; multiparous R2=0.068, p<0.0001) and **b**, pBMI category (primiparous, R2=0.041, p<0.0001; multiparous R2=0.071, p<0.0001). Significance was assessed by PERMANOVA blocked by sample time point as strata to acco unt for repeated measures. **c**, Bray-Curtis dissimilarity at each timepoint to all other timepoints within individual participants are negatively associated with pBMI in primiparous participants (n=40), but not in multiparous participants (n=25). Significance was analyzed within primiparous and multiparous participants by mixed linear model with pBMI, GWG category, and sample time point as interacting fixed effects and participant ID as a random effect.

### Parity modulates impact of pBMI and GWG on fecal SCFA levels

Because of their functional role in metabolic signalling pathways^19^, we assessed fecal SCFA level associations with pBMI and GWG across all timepoints in a subset of maternal fecal samples (primiparous, n=12; multiparous, n=11). This subset included samples that had enough starting material for SCFA quantification and was not biased in terms of pBMI, GWG, parity, maternal age, gestational length, or mode of delivery compared to participants not included in SCFA analyses. To account for variability in recovery between samples, the relative concentration of each SCFA was assessed as a percentage of total SCFAs (Fig. 2). Due to limited sample size, we assessed fecal SCFA levels by both GWG and pBMI—and the interaction of each of these factors with parity—across all samples.

**Figure 2.**
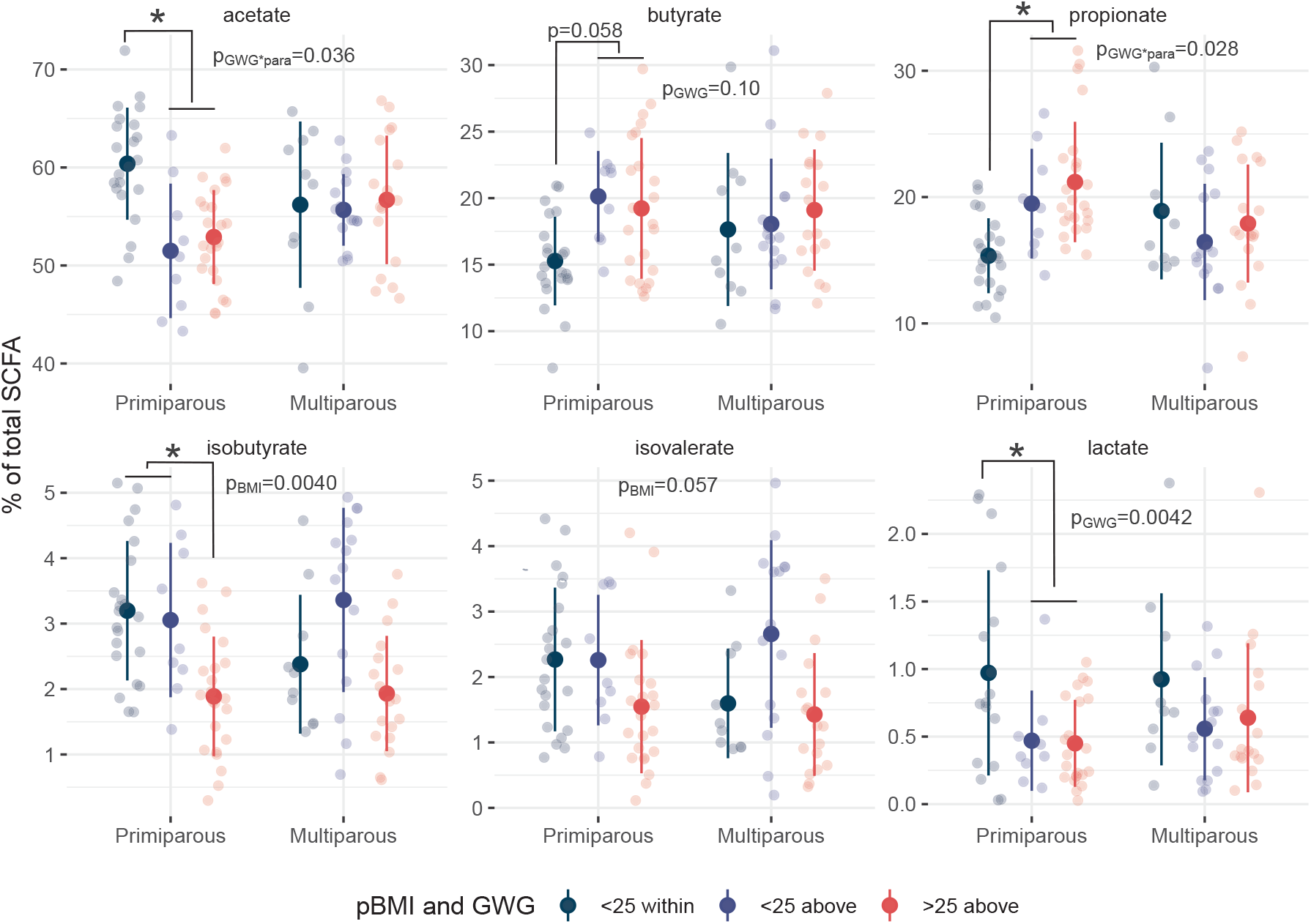
Relative SCFA concentrations differ with pBMI >25 and excess GWG. Dot plots of relative SCFA concentrations as a percentage of total SCFAs show decreased acetate (p=0.014) and lactate (p=0.034) with GWG above the recommended range and decreased isobutyrate with pBMI >25 (p=0.044). Primiparous (<25 within, n=7; <25 above, n=2; >25 above, n=7); Multiparous (<25 within, n=6; <25 above, n=6; >25 above, n=5). Individual points are shown (transparent dots) as well as the mean (solid black dot) and standard deviation (whiskers). Significance was assessed by mixed linear model with pBMI category or GWG category as fixed effects and participant ID as a random effect.

Fecal SCFA levels differed by GWG, and this effect was dependent upon parity as differences occurred only in primiparous participants (Supplementary Tables 1 and 2). Fecal acetate was decreased by GWG above the recommended range (main effect of GWG-by-parity interaction p=0.036) in primiparous participants (p=0.0041), and this decrease was accompanied by an increase in propionate (p=0.027, main effect of interaction p=0.028). Overall, lactate was decreased in participants with GWG above the recommended range (p=0.0042), particularly in primiparous participants (p=0.0047). Together, these data suggest that differences in non-digestible carbohydrate fermentation processes that produce SCFAs, and by inference the biochemical pathways between them, appear to be driven by gestational weight gain in primiparous women.

In contrast, differences in branched SCFA (BCFA) levels were driven by pBMI. Overall, isobutyrate was decreased by pBMI >25 (main effect of pBMI p=0.0040 vs pBMI <25), particularly in primiparous participants (p=0.013). Previous studies have shown BCFAs contribute to intestinal barrier function^20^ and facilitate the storage of diet-derived lipids in adipocytes by inhibiting lipolysis and increasing glucose uptake^21^. Thus, decreased BCFAs may contribute to a decrease in appropriate maternal metabolic adaptive flexibility leading to increased risk of pregnancy complications in participants with a pBMI >25 and excess GWG, though this hypothesis remains to be tested.

### Maternal taxonomic associations with pBMI, GWG, and parity

Taxa that were differentially abundant during pregnancy between pBMI and GWG categories were identified using DESeq2 (RRID: SCR_015687). Overall, 14 genera with a mean abundance > 150 differed by pBMI category (Fig. 3a-c, Supplementary Table 3) compared to participants with a pBMI < 25. Participants with a pBMI 25-30 had 6 differentially abundant genera, whereas participants with a pBMI > 30 had 11 genera that were differentially abundant (3 genera were differentially abundant for pBMI of both 25-30 and > 30). The relative abundance of *Blautia* was decreased in participants with a pBMI 25-30 or > 30 (Fig. 3a-c). This was accompanied by decreased *Bacteroides* and *Anaerostipes* in participants with pBMI >30. In contrast, *Prevotella 9* was markedly increased in participants with pBMI > 25 or > 30. In primiparous participants a pBMI of 25-30 or >30 was associated with increased *Rikenellaceae RC9 gut group*, which has been linked to higher propionate levels in high-fat fed mice^22^, consistent with the increase we observed in fecal propionate in participants with pBMI >25 and/or excess GWG.

**Figure 3.**
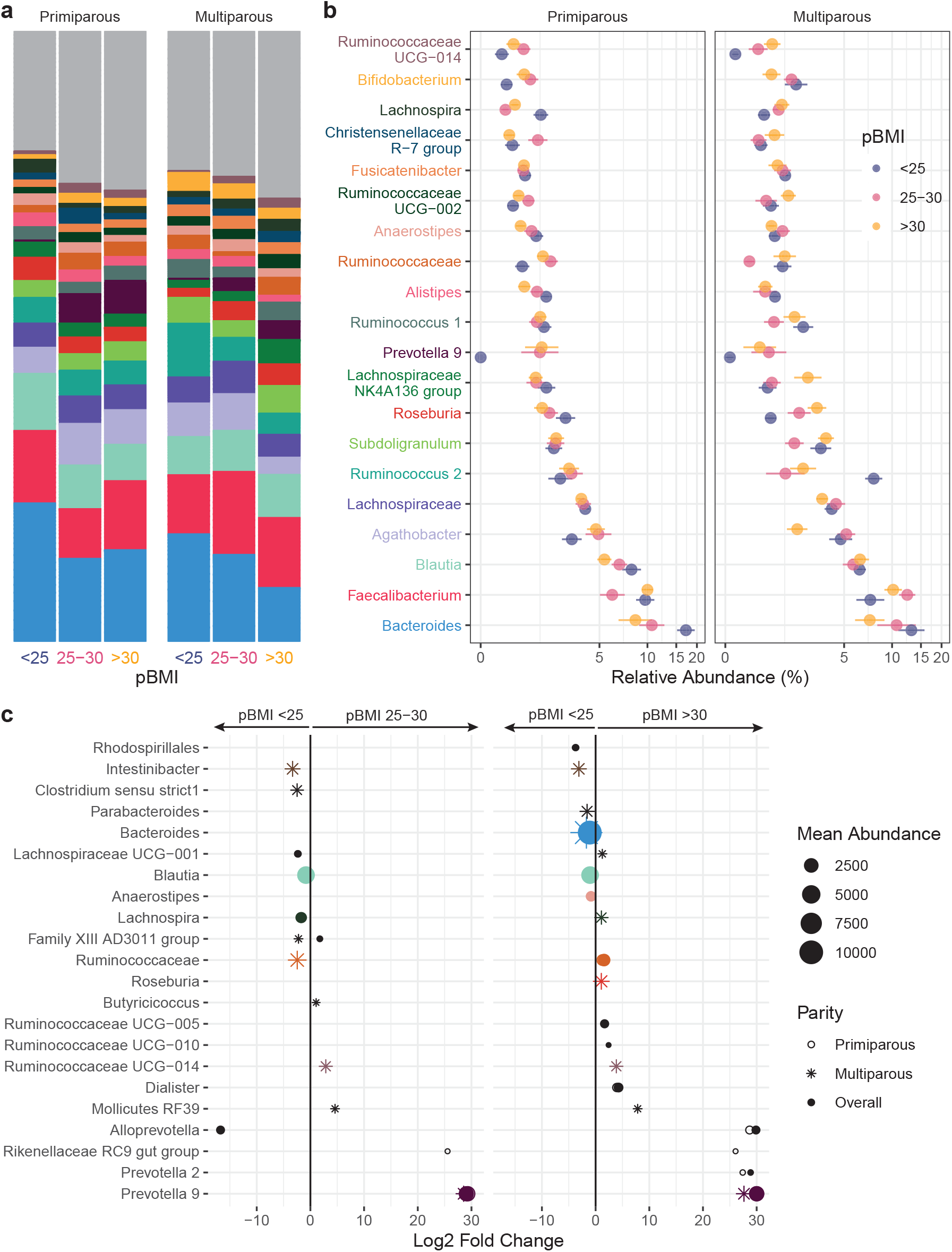
Taxonomic shifts with pBMI in pregnant gut microbiota. **a-b**, The relative abundances of the 20 most abundant genera in the maternal gut microbiota by pBMI <25 (blue, n=12), 25-30 (red, n=10), and >30 (yellow, n=12) are shown as a, mean relative abundance in taxa bar plots and b, mean ± standard error of the mean for primiparous (pBMI <25, n=7; 25-30, n=6; >30, n=7) and multiparous (pBMI <25, n=5; 25-30, n=4; >30, n=5) participant with excess GWG. **c**, Dot plots showing DESeq2 results of differentially abundant genera (mean abundance >150) by pBMI category overall (filled dots) and in primiparous (no fill dots) and multiparous (stars) participants (dot size based on mean abundance, colours correspond to genus colour in a where a differentially abundant genus is among the 20 most abundant genera).

The relative abundances of 13 genera with a mean abundance > 150 differed by GWG category (Fig. 4 a-c, Supplementary Table 3): compared to participants with GWG within the recommended range, participants with excess GWG (above) had 11 genera and participants with insufficient GWG (below) had 2 genera that were differentially abundant overall. *Prevotella 9* was markedly decreased by excess GWG, while insufficient GWG (below) was associated with decreased *Lachnospiraceae NK4A136 group*, the abundance of which has previously been associated with gut barrier function in mice^23^. Excess GWG was also associated with decreased relative abundances of 6 genera of the SCFA-producing family Ruminococcaceae (Fig. 4c), including uncultured genus (UCG) 005 which has previously been reported to decrease with gestational diabetes mellitus^24^. In primiparous participants excess GWG (above) was associated with a decreased in *Coprococcus 1*, which prior studies have found to be inversely associated with circulating triglyceride levels in non-pregnant individuals^25^. In multiparous participants, excess GWG was associated with increased *Bifidobacterium*. Overall, *Bifidobacterium* was also increased in multiparous participants compared to primiparous participants (Supplementary Table 3).

**Figure 4.**
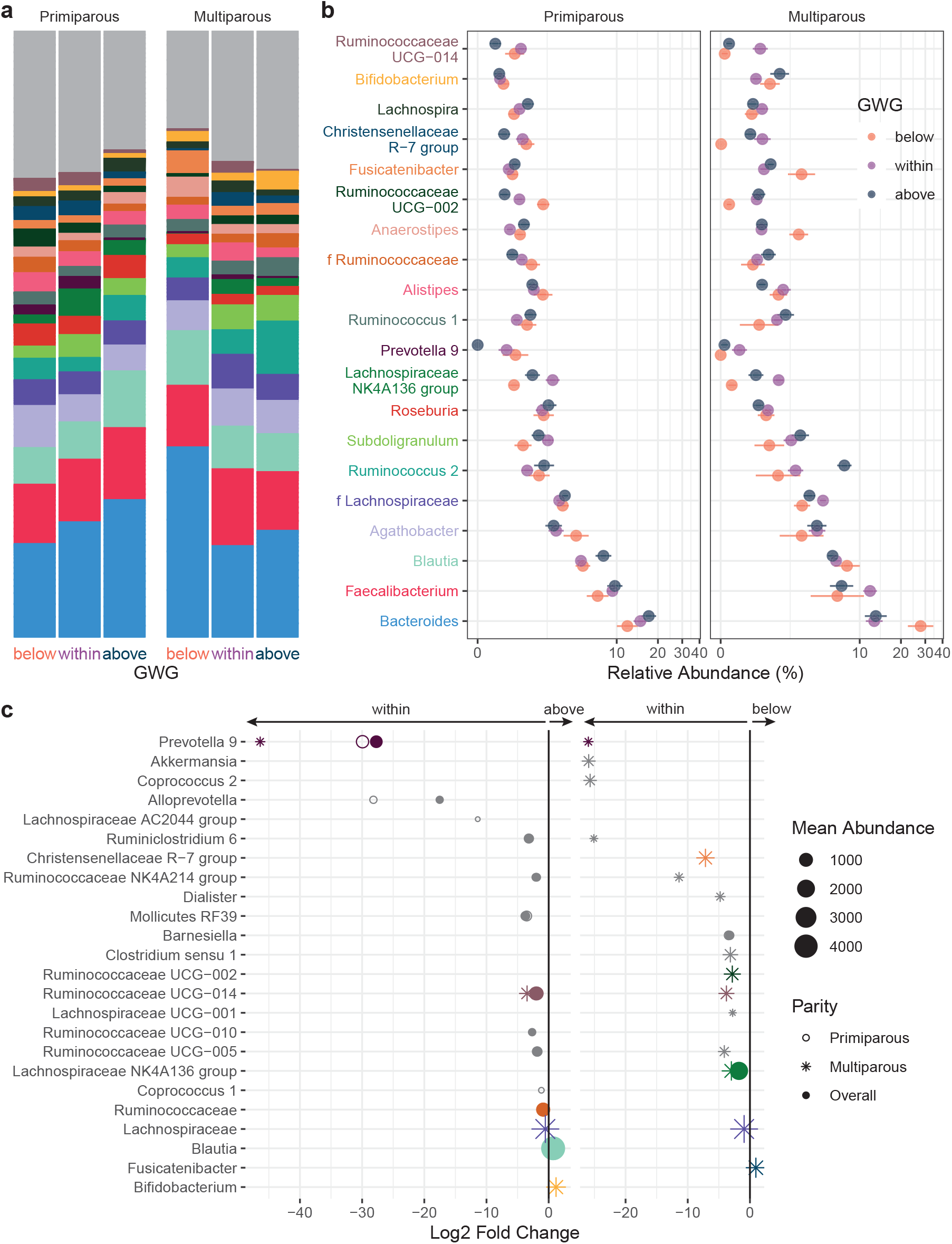
Taxonomic shifts with GWG in pregnant gut microbiota. **a-b**, The relative abundances of the 20 most abundant genera in the maternal gut microbiota by GWG below (orange, n=7), within (purple, n=17), and above (dark blue, n=12) the recommended range are shown as a, mean relative abundance in taxa bar plots and b, mean ± standard error of the mean for primiparous (below, n=5; within, n=11; above, n=7) and multiparous (below, n=2; within, n=6; above, n=5) participant with pBMI <25. **c**, Dot plots showing DESeq2 results of differentially abundant genera (mean abundance >150) by GWG category overall (filled dots) and in primiparous (no fill dots) and multiparous (stars) participants (dot size based on mean abundance, colours correspond to genus colour in a where a differentially abundant genus is among the 20 most abundant genera).

### Maternal GWG and the infant gut microbiota at 6 months of age

Of the 63 total mother-infant dyads who provided an infant fecal sample at 6 months of age, we excluded 3 for nicotine exposure during pregnancy, 1 for pre-term birth, and 1 for low sequencing read count, which left 58 infants for full analysis. The majority of infants delivered by C-section were exposed to perinatal antibiotics (12/14; 86%;) while relatively few vaginally delivered infants were exposed (9/44; 20%). As perinatal antibiotic exposure has previously been shown to disturb vertical (maternal-to-infant) transmission of the microbiota during birth^26^, we assessed the impact of perinatal antibiotics (between 33 weeks gestation and birth, including antibiotics given during delivery) on the infant gut microbiota in this cohort. There was no significant impact of antibiotics on alpha diversity (Observed p=0.70, Shannon p=0.39; Extended Data Fig. 3a). Infant gut microbiota beta-diversity differed by antibiotic exposure in those born to primiparous participants (PERMANOVA, R^2^=0.068, p=0.019), but not in those born to multiparous participants (R^2^=0.043, p=0.56; Extended Data Fig. 3b). This may reflect the a rescue effect due to horizontal transmission from older siblings^27^. Overall, perinatal antibiotic exposure was associated with a decreased abundance of 8 genera, including *Bifidobacterium, Streptococcus, Blautia*, and *Parabacteroides* (Extended Data Fig. 4 a-c). Three additional genera (*Lachnoclostridium, Flavonifractor*, *Intestinibacter*) were significantly decreased by antibiotic exposure only in primiparous participants. *Akkermansia* and an unidentified genus of the family Lachnospiraceae were the only genera that increased with antibiotic exposure, although only in primiparous participants. In contrast, both these genera were decreased by antibiotic exposure in multiparous participants. These data are consistent with others that show that perinatal antibiotic exposure significantly perturbs the infant gut microbiota^26^ but extend these observations to show that parity influences these effects.

In infants that were not exposed to antibiotics (female, n=19; male, n=18) the majority were delivered vaginally (35/37; 95%). Infant characteristics at birth did not differ by maternal pBMI category overall or within primiparous and multiparous participants (Extended Data Table 2). Although infant birth weight did not differ by GWG overall, excess GWG was associated with higher birth weight in multiparous participants (p=0.028; Extended Data Table 2); although this difference was not significant after adjusting for multiple testing. Overall, at 6 months of age, infant weight did not differ significantly by maternal pBMI, GWG, or parity. Among infants with available feeding data, at 6 months of age the majority were both breastfed (27/30; 90%) and eating solid foods (23/28; 82%).

We evaluated the impacts of maternal pBMI and GWG on the gut microbiota of infants at 6 months of age. We identified a total of 1063 ASVs after removal of host-associated reads (Kingdom Eukaryota, Order Chloroplast, Family Mitochondria, or no assigned Phylum). The median sample read count was 59,561 (minimum 6,045; maximum 124,303). Overall, infant microbiota alpha diversity (number of observed ASVs) differed by maternal pBMI (p = 0.036) and GWG category (p = 0.047; Fig. 5a) but did not differ by infant sex (p=0.78). There was a significant effect of maternal GWG on beta-diversity in infants of multiparous (R^2^=0.378, p=0.0009), but not primiparous, mothers (R^2^=0.0950, p=0.377; Fig. 5b). Infant beta-diversity did not differ by maternal pBMI category (R^2^=0.0511, p=0.565) or infant sex (R^2^ = 0.0126, p=0.914). Therefore, in addition to previous reports linking maternal GWG and BMI to shifts in the infant gut microbiota^13,28–33^, we now show that that parity modulates the impact of maternal GWG on the infant gut microbiota.

**Figure 5.**
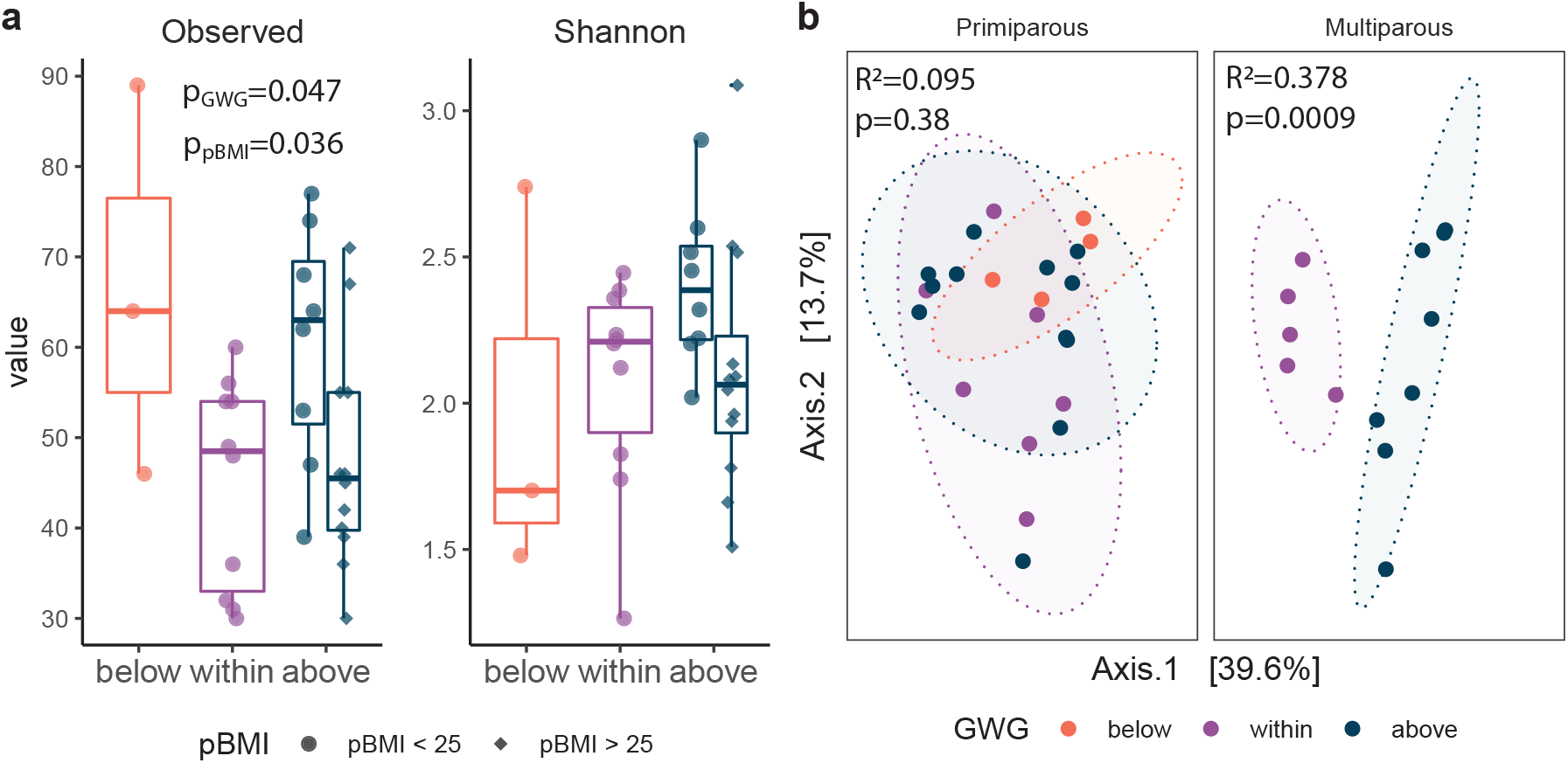
Parity modulates impact of GWG on infant gut microbiota. **a**, Box plots of alpha diversity by gestational weight gain (GWG) category in primiparous participants (n=24) and multiparous participants (n=13). Alpha diversity (observed ASVs) was significantly increased by GWG above the recommended range (p=0.034) in infants of participants with pBMI <25. Significance assessed by a linear model (Kenward–Roger d.f.), with GWG category as a fixed effect. The box plot centre line represents the median; the box limits represent the upper and lower quartiles; the whiskers represent the 1.5× interquartile range; the points represent the outliers. **b**, PCoA of Bray-Curtis distances shows modest clustering by GWG category (primiparous, R2=0.039, p<0.0001; multiparous R2=0.068, p<0.0001). Primiparous (below, n=4; within, n=7; above=12); Multiparous (within, n=5; above, n=8). Significance was assessed by PERMANOVA.

Taxa that were differentially abundant in infants between maternal pBMI and GWG categories were identified using DESeq2. Overall, maternal excess GWG (above) was associated with an increased abundance of *Bifidobacterium* (Fig. 6 a-c)—echoing similar shifts in the maternal gut microbiota (Fig. 4 a-c). In infants of multiparous, but not primiparous, participants this was accompanied by a decrease in *Bacteroides*. Compared to a maternal pBMI 18.5-25, a maternal pBMI >25 was associated with a decreased relative abundance of Clostridiales genera *Blautia* and *Intestinibacter* in the infant gut microbiota, which were also decreased by pBMI >25 in the maternal gut microbiota. Depleted *Blautia* in obese children has been linked to intestinal inflammation^34^ which is consistent with an observed association between excess maternal adiposity with childhood obesity and inflammation^35^. This overlap in differentially abundant taxa between the maternal and infant gut microbiota is consistent with vertical transmission of the maternal gut microbiota^36^.

**Figure 6.**
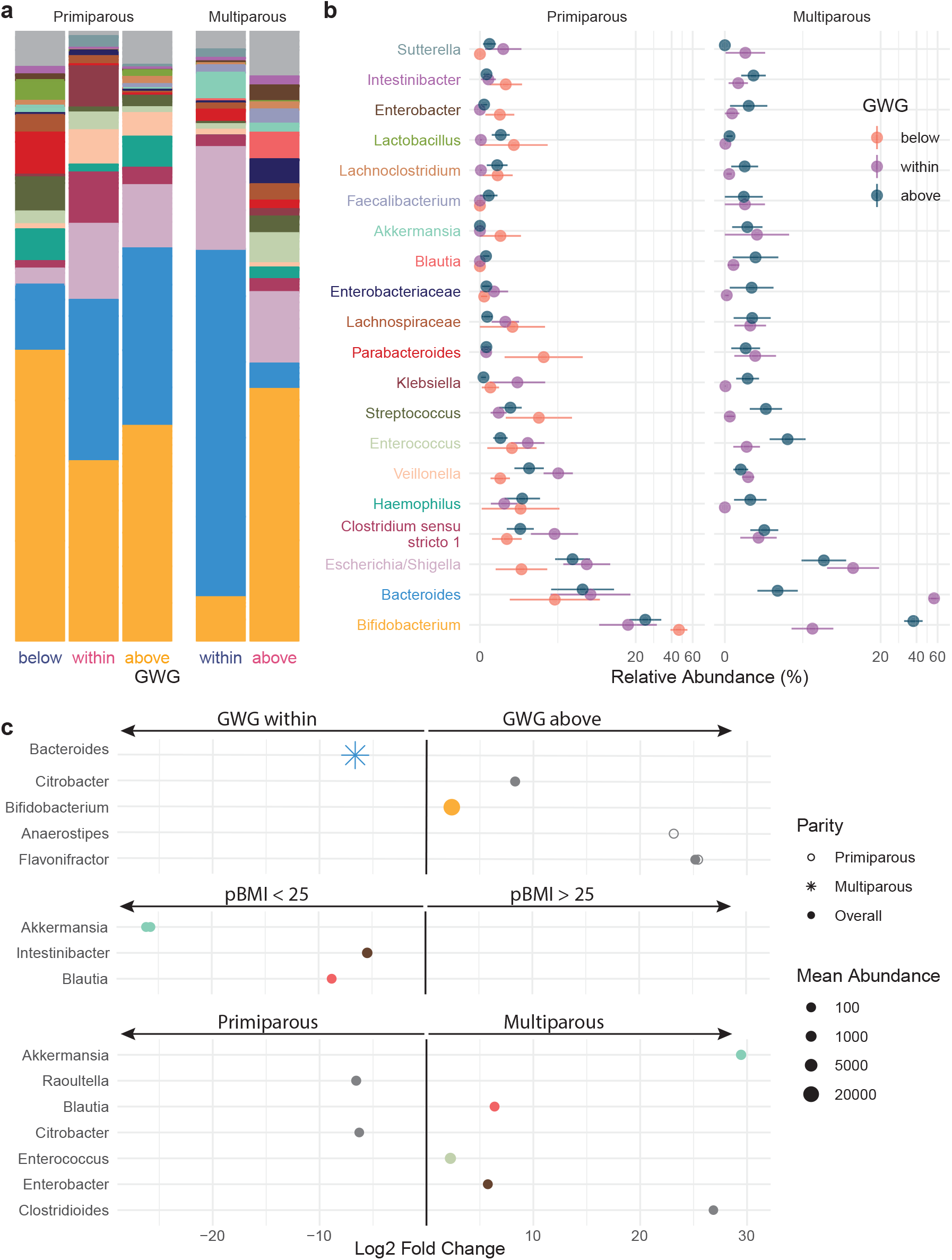
Taxonomic shifts with maternal GWG in infant gut microbiota. **a-b**, The relative abundances of the 20 most abundant genera in the infant gut microbiota by GWG below (orange), within (purple), and above (dark blue) the recommended range are shown as a, mean relative abundance in taxa bar plots and b, mean ± standard error of the mean for infants of primiparous (below, n=4; within, n=7; above, n=12) and multiparous (within, n=5; above, n=8) participants. **c**, Dot plots showing DESeq2 results of differentially abundant genera overall (filled dots) and in primiparous (no fill dots) and multiparous (stars) participants by maternal GWG, BMI, and parity. Dot size based on mean abundance, colours correspond to genus colour in a where a differentially abundant genus is among the 20 most abundant genera.

## Discussion

Pregnancy-associated shifts in maternal gut microbiota have been implicated in mediating metabolic adaptations to pregnancy^4^, but few studies have investigated the gut microbiota longitudinally over the course of healthy pregnancy. Because of this, there exists contrasting data regarding whether these temporal shifts do^4^ or do not^9,37^ occur. In this study, we investigated the independent and combined impacts of maternal pre-pregnancy BMI (pBMI) and gestational weight gain (GWG) on the maternal gut microbiota over the course of pregnancy and lactation, and infant gut microbiota at 6 months of age. We show pregnancy-associated remodeling of the gut microbiota is individual-specific and therefore longitudinal sampling within individuals is essential. We found that pBMI and GWG impact the maternal gut microbiome during pregnancy—both the microbial composition and possible function via SCFA metabolites—and also impact the infant gut microbiota at 6 months of age. We also show for the first time in humans that parity modulates the impact of pBMI on pregnancy-associated remodeling of the maternal gut microbiota and also modulates the impact of perinatal antibiotic exposure and GWG on the infant gut microbiota. We now suggest that the maternal gut microbiota may retain a “ecological memory”^38^ of previous pregnancies.

We found that in primiparous (first) pregnancy, the gut microbiota of participants with higher pBMI changed less over the course of pregnancy and postpartum than that of participants with a lower pBMI. Previous studies of the gut microbiota have found little-to-no overall effect of pBMI on overall beta-diversity in the first trimester^39^ and across pregnancy^8,37^ but have not investigated beta-diversity within individuals over the course of pregnancy. Shifts in the gut microbiome due to diet or antibiotic perturbation are highly individual-specific^40^ and our data now suggest pregnancy-associated remodeling of the gut microbiota is also individual-specific. Our observed decrease in intra-individual beta-diversity with increasing pBMI in primiparous participants is consistent with a decreased magnitude of metabolic adaptation to pregnancy with increasing pBMI^41–43^, and highlights the necessity of longitudinal sampling for investigations of pregnancy-associated microbiota remodeling.

Our novel observation that this remodeling was not associated with pBMI in multiparous participants suggests that microbiotal adaptations to first pregnancy may have persistent impacts that modulate microbiotal adaptations to subsequent pregnancies. The impact of parity on the gut microbiome and its contribution to metabolic adaptations to pregnancy has not previously been investigated in human pregnancy. Recently, Berry et al.^44^ found that maternal microbiotal adaptations to pregnancy occurred more rapidly with increasing parity in pigs. This may reflect increased adaptive plasticity resulting from adaptations to first pregnancy^45^. Indeed, it is known that first-pregnancies are associated with greater maternal constraint^46,47^ compared to subsequent pregnancies, which may include reduced or delayed metabolic adaptations to pregnancy.^48^ Whether a functional ecological memory exists from one pregnancy to the next in humans is still unclear, but our data support the notion that parity influences microbial community composition and must be considered when investigating human pregnancy-related microbiotal (and intestinal) adaptations.

Maternal adaptations to pregnancy include increased intestinal surface area and increased transit time, allowing increased nutrient uptake to provide the energy required by fetal development and lactation^49^. SCFAs may support these intestinal adaptations by promoting proliferation of the intestinal epithelium^50^ and improved intestinal barrier funciton^20,51^. Although human data on intestinal adaptations to pregnancy are limited, in animal models we have found that excess adiposity during pregnancy is associated with decreased intestinal length and increased intestinal permeability^52^. In this study, we found that parity modulated the impact of pBMI and GWG on the relative abundance of maternal fecal SCFAs. In primiparous participants, GWG above the recommended range was associated with decreased acetate and increased propionate. As fecal SCFAs levels may be more strongly correlated to intestinal absorption than production^53^, these changes could be due to altered epithelial cell uptake resulting from reduced intestinal adaptations to pregnancy, but this hypothesis remains to be tested.

Alternatively, differences in fecal SCFA levels may result from shifts in microbial production due to altered microbiota community composition. We found increased propionate in primiparous participants with excess GWG and/or pBMI>25, consistent with a prior study that found increased fecal propionate in pregnant individuals with a pBMI>25 compared to pBMI<25, although parity and GWG were not reported^54^. *Prevotella*-dominant microbiota produce more propionate—and more SCFAs overall—than *Bacteroides*-dominant microbiota^55^. This is consistent with our finding that *Prevotella* was increased in participants with a pBMI>25, while *Bacteroides* was decreased, particularly in primiparous participants. A previous study also found decreased *Bacteroides* in the first and third trimester with pre-pregnancy overweight^11^, although others have reported increased levels in the third trimester with pre-pregnancy overweight but not obesity^12,56^. In independent studies, *Prevotella* has been associated with improved glucose tolerance^57^ and impaired glucose tolerance^58^. This functional diversity mirrors that of *Bacteroides* and suggests the impacts of these genera depend on host physiology including pregnancy history. Future studies are needed to longitudinally investigate pregnancies within participants as well as in the intrapartum period, to test this hypothesis.

Shifts in the maternal gut microbiota can impact the infant gut microbiome by (1) altering fetal intestinal development through the *in-utero* environment and (2) direct transmission of maternal microbes to the infant during and after birth. Disruption of this vertical transmission may have persistent effects on the infant gut microbiota, as we found perinatal antibiotic exposure was associated with a decrease in *Bifidobacterium* in the infant gut microbiota at 6 months of age. We show novel data, that supports the position that parity modulates the impact of maternal GWG on infant characteristics at birth and the infant gut microbiota at 6 months of age. Gestational weight gain above the recommended range is often associated with a reduced risk of small for gestational age (SGA) birth weight and an increased risk of large for gestational age (LGA) birth weight^59^, although some data suggest that maternal obesity can also be associated with intrauterine growth restriction (IUGR)^60^. Little to no data exist on the modulating effect of parity. We found excess GWG was associated with higher birth weight in infants of multiparous participants, and a tendency towards lower birthweight in infants of primiparous participants. These impacts of excess GWG were also associated with a shift in microbiota beta-diversity at 6 months of age in infants of multiparous—but not primiparous—participants. These same infants also had a decrease in *Bacteroides* and increase in *Bifidobacterium* reflecting similar shifts in the maternal gut microbiota consistent with the high rates of vertical transmission previously found for these genera^61^. Therefore, health behaviours during pregnancy and the postpartum/intrapartum interval should be supported not only for maternal health but also infant microbiotal health.

Our study has both strengths and limitations. Our longitudinal sampling of a prospective pregnancy cohort allowed us to explore beta-diversity within participants to overcome the overwhelming effect of inter-individual variation. However, as we recruited participants in the first trimester of pregnancy, we are not able to address pregnancy-associated shifts in the gut microbiota that may occur very early in gestation. We also show potential functional outcomes of observed microbial shifts in our analysis of fecal SCFA levels, but did not quantify circulating SCFA levels, which may more directly modulate host metabolism^62^. We also cannot measure whether production versus absorption on these SCFAs is altered with pregnancy, GWG, BMI or parity. Finally, although our study is unique among studies of the pregnancy microbiome in its design and sample size and relationship to parity, we were unable to examine any interacting effects of pBMI and GWG as the vast majority of participants with high pBMI also had GWG above the recommended range.

In conclusion we show that parity modulates impact of pBMI on pregnancy-associated remodeling of the maternal gut microbiota, colonization of the neonatal gut, and the impact of GWG on the infant gut microbiota. We also show that in women in their first pregnancy, high pBMI is associated with diminished remodeling of the gut microbiome during pregnancy, which may reflect a reduced requirement for metabolic adaptations to pregnancy. This is accompanied by functional impacts via altered SCFA levels, which may reflect reduced intestinal adaptations to pregnancy. These outcomes are important not only to maternal health, but also to infant health as infants born to multiparous women with excess gestational weight gain show distinct shifts in their microbiota. Finally, our data suggest the human gut microbiome may retain an “ecological memory”^38^ of prior pregnancies^44^ which may serve as an adaptation that minimizes maternal constraint in subsequent pregnancies.Acknowledgements

We thank all the participants that were recruited in this study. We would like to thank Alexandra Kühn, Dr. Hanna Brinkmann, Laura Pasura, Laura Maschirow, Alexander Schwickert, and Sonja Entringer for assisting with patient recruitments; Loreen Ehrlich, Kerstin Melchior, and Thomas Ziska for assisting with sample preparation. We thank Michelle Shah for performing genomic DNA extractions and Laura Rossi for preparing amplicons for sequencing. M.G.S. and D.M.S. are supported by the Canada Research Chairs Program. T.B. is supported by Deutsche Forschungsgemeinschaft grant no. BR2925/10-1. A.P. is supported by Deutsche Forschungsgemeinschaft grant no. PL241/16-1.

## Author Contributions

Conceptualization, K.M.K., S.A., T.B., A.P., and D.M.S.; Investigation, K.M.K, J.S., M.H., and T.B.; Data Curation, K.M.K; Formal Analysis, K.M.K.; Visualization, K.M.K; Writing – Original Draft, K.M.K; Writing – Review & Editing, K.M.K, M.G.S., T.B., and D.M.S; Supervision, J.F.R.B., T.B., A.P., W.H., and D.M.S.; Project Administration, T.B., A.P., W.H., and D.M.S.; Resources, T.B., A.P., W.H., M.G.S., and D.M.S.; Funding Acquisition, T.B. and D.M.S.

## Competing Interests statement

The authors declare that they have no competing interests.

## Methods

### Data and code availability

Sequencing data have been deposited at the NCBI Sequence Read Archive (SRA; PRJNA878704) and are publicly available as of the data of publication. All original code has been deposited on GitHub and is publicly available as of the date of publication. Any additional information required to reanalyze the data reported in this paper is available from the lead contact upon request.

### Study design

The study protocol was reviewed and approved by the Charité ethics committee (EA 4/059/16). All participants provided written informed consent. Study participants were part of an ongoing research program at the Charité University Berlin. Healthy females >18 years of age with singleton pregnancies were prospectively recruited between 9-17 weeks of gestation. We excluded participants who (1) had severe chronic gastrointestinal diseases or conditions, (2) had any significant heart, kidney, liver, or pancreatic diseases, (3) had pre-existing diabetes, (4) had depression, or (5) delivered infants with severe malformations noted at birth. A total of 89 participants were included in the study. We additionally excluded maternal and infant samples from participants who smoked during pregnancy and excluded maternal samples from participants who took antibiotics during pregnancy. After these exclusions, this study included infant samples from 58 participants, and maternal samples from 65 participants.

### Sample collection

Participants were evaluated at the end of the first trimester (9-17 weeks gestation), second trimester (26-28 weeks), and third trimester (>33 weeks) of pregnancy, at delivery, and at 6 months postpartum. Prepregnancy body mass index (pBMI) was obtained from measured height recruitment and recalled prepregnancy weight. BMI was categorized into normal weight (18.5-24.9kg/m2), overweight (25-29.9 kg/m2), and obese (>30<40 kg/m2). At each study visit, participant weight was measured. Participant collected fecal samples at home prior to each study visit following a standardized sterile stool collection protocol. Stool was frozen at −20°C immediately after collection in provided sample collection bags and stored at −80°C upon arrival until further use. None had taken prebiotics, laxatives, or diarrhea inhibitors in the days before the sampling.

### DNA extraction and amplification

Genomic DNA (gDNA) was extracted from fecal samples as described previously^63^ on a MagMAX Express semi-automatic robot (MagMAX-96 DNA Multi-Sample Kit; Invitrogen, cat# 4413022) with the addition of a mechanical lysis step using 0.2 g of 2.8 mm ceramic beads to improve extraction efficiency and without mutanolysin. Four negative controls were included on each extraction plate. PCR amplification of the variable 3 and 4 (V3-V4) regions of the 16S rRNA gene was performed on the extracted DNA using methods previously described^64^. Each reaction contained 5 pmol of primer (341F – CCTACGGGNGGCWGCAG, 806R – GGACTACNVGGGTWTC-TAAT), 200 mM of dNTPs, 1.5μl 50 mM MgCl2, 2 μl of 10 mg/ml bovine serum albumin (irradiated with a transilluminator to eliminate contaminating DNA) and 0.25μl Taq polymerase (Life Technologies, Canada) for a total reaction volume of 50 μl. 341F and 806R rRNA gene primers were modified to include adapter sequences specific to the Illumina technology and 6-base pair barcodes were used to allow multiplexing of samples. Lack of amplification in all negative controls was confirmed by gel electrophoresis and one negative control from each extraction plate was sequenced.

### 16S rRNA gene sequencing

16S DNA products of PCR amplification were sequenced using the Illumina MiSeq platform (2×300bp) at the Farncombe Genomics Facility (McMaster University, Hamilton ON, Canada). Primers were trimmed from FASTQ files using Cutadapt^65^ (RRID: SCR_011841) and DADA2^66^ was used to derive amplicon sequence variants (ASVs). Taxonomy was assigned using the Silva 132 reference database^67^. Non-bacterial ASVs were culled (kingdom Eukaryota, family Mitochondria, order Chloroplast, or no assigned phylum), as was any ASV to which only 1 sequence was assigned.

### Quantification of short-chain fatty acids

SCFA levels were measured in a subset of fecal samples by gas-chromatography mass spectroscopy (GC-MS) at the McMaster Regional Centre of Mass Spectrometry. A weight equivalent amount of 3.7% HCl, 10 μL of internal standard, and 500 μL of diethyl ether was added to each fecal sample and vortexed for 15 min. After vortexing, 400 μL of diethyl ether fecal extract was transferred to a clean 1.5 mL Eppendorf tube. In a chromatographic vial containing an insert, 20 μL of N-tert-butyldimethylsilyl-N-methyltrifluoroacetamide was added, after which 60 μL of diethyl ether fecal extract was added. The mixture was incubated at room temperature for 1 hour and analyzed using gas chromatography-mass spectrometry (6890N GC, coupled to 5873N Mass Selective Detector; Agilent Technologies, Santa Clara, CA, USA). Statistical significance was assessed by mixed linear model with pBMI, GWG category, and parity as fixed effects and participant ID as a random effect, and multiple comparisons by pBMI and GWG category were performed within primiparous and multiparous participants (Kenward-Rogers degrees of freedom with Tukey adjustment for multiple comparisons).

### Statistical analysis

#### Participant characteristics

Participant characteristic variables (Extended Data Tables 1 and 2) were analyzed in R using the gtsummary package (RRID: SCR_021319). Normality was determined by Shapiro-Wilk test (rstatix; RRID: SCR_021240) and significance was assessed by Kruskal-Wallis test and false discovery rate correction for multiple testing.

#### 16S analysis

We performed alpha and beta-diversity analyses in R using phyloseq^68^ (RRID: SCR_013080). Alpha diversity analyses were performed on rarefied data (14872 reads per sample). Significance of alpha diversity was analyzed by linear mixed model (Kenward–Roger degrees of freedom) with time point, pBMI, or GWG category as a fixed effect and participant ID as a random effect. Beta-diversity analyses were performed on proportionally normalized data. We tested for whole community differences across groups using vegan’s^69^ (RRID: SCR_011950) implementation of permutational multivariate analysis of variance (PERMANOVA) in the adonis2 command blocked by sample time point as strata to account for repeated measures. These results were visualized via Principal Coordinate Analysis (PCoA) ordination using R’s ggplot2 package (RRID: SCR_014601)^70^. Significance of Bray-Curtis dissimilarity was analyzed within primiparous and multiparous participants by mixed linear model with pBMI, GWG category, and sample time point as interacting fixed effects and participant ID as a random effect. Differential abundance analysis was performed using DESeq2 (RRID: SCR_015687) across all pregnancy time points. We did not assess interacting effects of pBMI and GWG as (1) pBMI category and GWG category were collinear (χ-squared p < 0.0001) and (2) all but 4 participants with a pBMI > 25 experienced excess GWG. These 4 participants were excluded and the impact of pBMI was assessed within participants with excess GWG and the impact of GWG was assessed in participants with a pBMI 18.5-25.

#### SCFA analysis

Statistical significance was assessed by mixed linear models with pBMI category and parity or GWG category and parity as fixed effects and participant ID as a random effect, and multiple comparisons by pBMI and GWG category were performed within primiparous and multiparous participants (Kenward-Rogers degrees of freedom with Tukey adjustment for multiple comparisons). All statistical details, including statistical tests used, sample sizes, and definition of center and dispersion can be found in the figure legends.

---

**Extended Data Figure 1.**
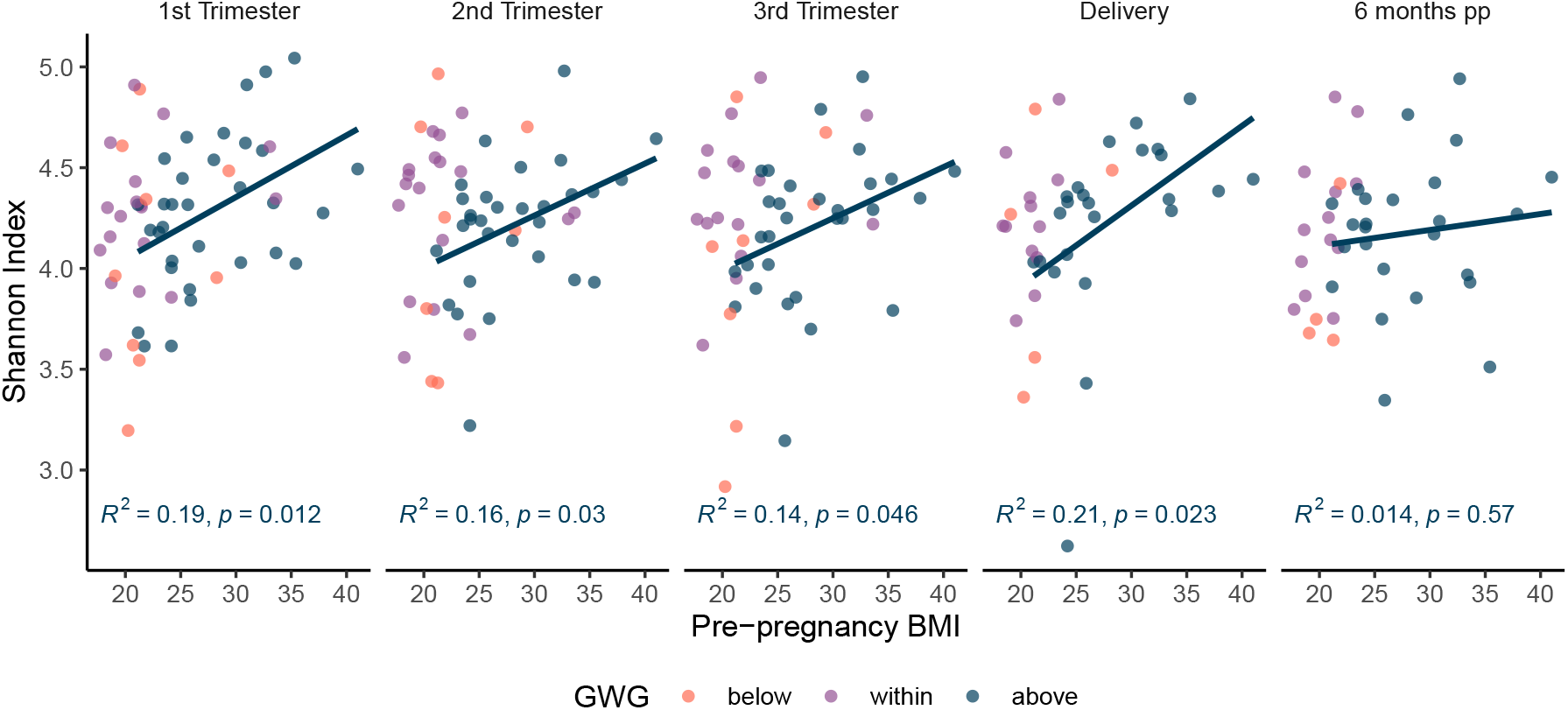
Alpha diversity correlated with pBMI during pregnancy. Boxplots of Shannon alpha diversity overlayed with dot plots show a positive correlation with pBMI at each pregnancy timepoint (p=0.012-0.046) but not at 6 months postpartum (p=0.57) in participants with excess GWG and no overall difference in alpha diversity between timepoints across all participants. Significance assessed by linear model (Kenward–Roger d.f.), with pBMI as a fixed effect. The box plot centre line represents the median; the box limits represent the upper and lower quartiles; the whiskers represent the 1.5× interquartile range; the points represent the outliers.

**Extended Data Figure 2.**
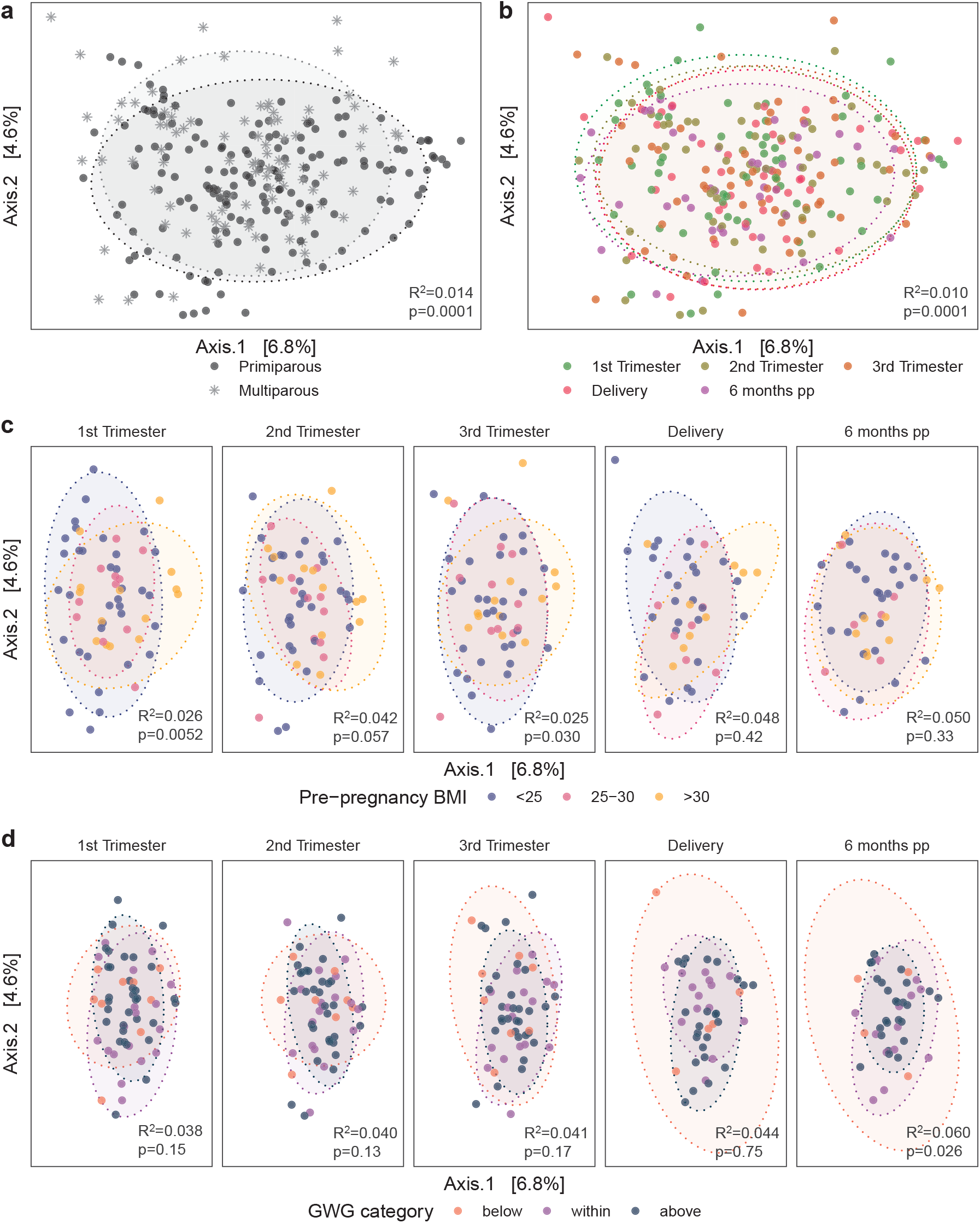
Impacts of pBMI, GWG, parity, and sample timepoint on maternal gut microbiome beta diversity. PCoA of Bray-Curtis distances shows modest clustering by **a**, parity (R^2^=0.014, p<0.0001) and **b**, timepoint (R^2^=0.010, p<0.0001). **c**, Clustering by pBMI category within timepoints is significant in the first trimester (R^2^ =0.026, p=0.0052) and the third trimester (R^2^ =0.025, p=0.033). **d**, Clustering by GWG category was significant at 6 months post-partum (R^2^=0.060, p=0.026). First trimester, n=62; second trimester, n=58; third trimester, n=56; delivery, n=41; 6 months post-partum, n=43. Significance was assessed by PERMANOVA.

**Extended Data Figure 3.**
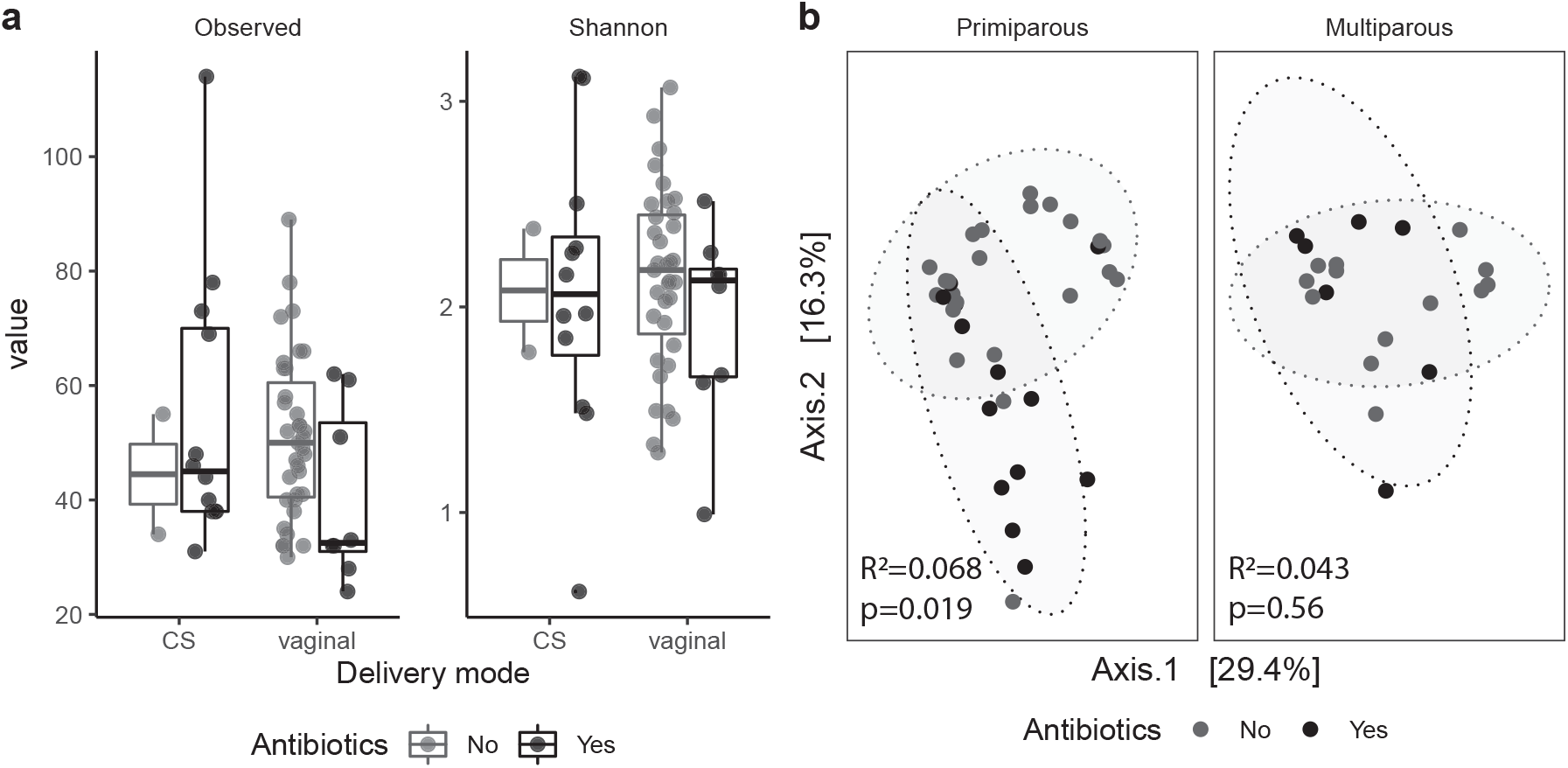
Parity modulates impact of peripartum antibiotics on infant gut microbiota. **a,** Box plots of alpha diversity at 6 months of age shows no effect of peripartum antibiotic exposure. Significance assessed by a linear model (Kenward–Roger d.f.), with antibiotic exposure as a fixed effect. The box plot centre line represents the median; the box limits represent the upper and lower quartiles; the whiskers represent the 1.5× interquartile range; the points represent the outliers. **b,** PCoA of Bray-Curtis distances in primiparous (n=38) and multiparous (n=20) participants shows modest clustering by GWG category (primiparous, R2=0.039, p<0.0001; multiparous R2=0.068, p<0.0001). Significance was assessed by PERMANOVA.

**Extended Data Figure 4.**
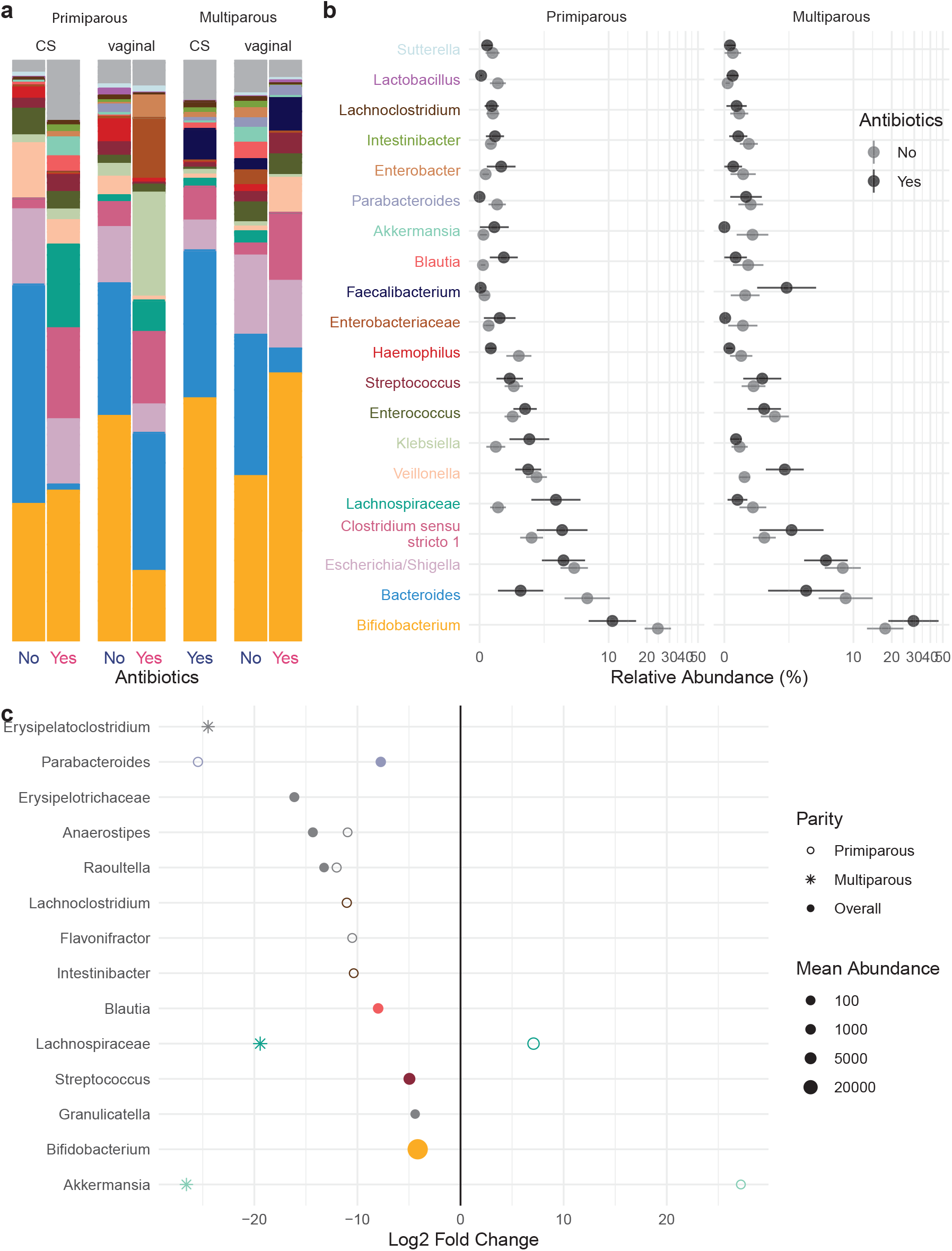
Taxonomic shifts with peripartum antibiotics in infant gut microbiota. **a-b,** The relative abundances of the 20 most abundant genera in the infant gut microbiota by peripartum antibiotic exposure are shown as a, mean relative abundance in taxa bar plots and b, mean ± standard error of the mean for primiparous (n=38) and multiparous (n=20) participant with pBMI <25. **c**, Dot plots showing DESeq2 results of differentially abundant genera by perinatal antibiotic exposure (dot size based on mean abundance, colours correspond to genus colour in a where a differentially abundant genus is among the 20 most abundant genera).

**Extended Data Table 1.** Participant characteristics.

| Characteristic | Parity |  |  |  | Overall pBMI |  |  |  |  | Primiparous pBMI |  |  |  |  | Multiparous pBMI |  |  |  |  |
| --- | --- | --- | --- | --- | --- | --- | --- | --- | --- | --- | --- | --- | --- | --- | --- | --- | --- | --- | --- |
|  | primiparous<sup>†</sup> | multiparous<sup>†</sup> | p<sup>2</sup> | q<sup>3</sup> | <25<sup>†</sup> | 25-30<sup>†</sup> | >30<sup>†</sup> | p<sup>2</sup> | q<sup>3</sup> | <25<sup>†</sup> | 25-30<sup>†</sup> | >30<sup>†</sup> | p<sup>2</sup> | q<sup>3</sup> | <25<sup>†</sup> | 25-30<sup>†</sup> | >30<sup>†</sup> | p<sup>2</sup> | q<sup>3</sup> |
| n | 40 | 25 |  |  | 39 | 12 | 14 |  |  | 24 | 7 | 9 |  |  | 15 | 5 | 5 |  |  |
| Age | 29.6 (4.4) | 32.8 (3.1) | 0.002 | **0.009** | 30.9 (4.3) | 31.3 (4.6) | 30.3 (3.9) | 0.8 | 0.8 | 29.9 (4.6) | 29.6 (5.3) | 29.1 (3.5) | >0.9 | >0.9 | 32.60 (3.20) | 33.80 (2.17) | 32.40 (4.04) | 0.8 | >0.9 |
| Primigravida | 27 (68%) | 0 (0%) | <0.001 | **<0.001** | 17 (44%) | 3 (25%) | 7 (50%) | 0.4 | 0.8 | 17 (71%) | 3 (43%) | 7 (78%) | 0.3 | 0.6 | 0 (0%) | 0 (0%) | 0 (0%) |  |  |
| pBMI | 23.8 (21.0, 29.0) | 23.5 (21.2, 26.6) | >0.9 | >0.9 | 21.3 (19.6, 23.2) | 26.4 (25.8, 28.4) | 33.2 (31.3, 34.9) | <0.001 | **<0.001** | 21.2 (19.4, 22.0) | 28.0 (25.7, 28.8) | 33.4 (32.7, 33.6) | <0.001 | **<0.001** | 21.3 (20.2, 23.4) | 26.1 (25.9, 26.6) | 31.0 (30.9, 35.3) | <0.001 | **<0.001** |
| GWG (kg) | 15.0 (12.0, 19.0) | 15.0 (13.0, 20.0) | 0.8 | >0.9 | 15.0 (13.0, 18.3) | 16.5 (14.6, 21.1) | 13.5 (10.0, 19.4) | 0.6 | 0.8 | 14.9 (13.0, 17.3) | 19.0 (15.5, 21.8) | 12.0 (10.0, 18.0) | 0.4 | 0.6 | 15.0 (13.0, 20.0) | 15.0 (14.8, 16.0) | 16.0 (10.0, 19.9) | >0.9 | >0.9 |
| GWG category |  |  | 0.5 | 0.8 |  |  |  | 0.001 | **0.004** |  |  |  | 0.014 | 0.056 |  |  |  | 0.069 | 0.2 |
| below | 6 (15%) | 3 (13%) |  |  | 7 (19%) | 2 (17%) | 0 (0%) |  |  | 5 (22%) | 1 (14%) | 0 (0%) |  |  | 2 (15%) | 1 (20%) | 0 (0%) |  |  |
| within | 13 (33%) | 6 (26%) |  |  | 17 (47%) | 0 (0%) | 2 (14%) |  |  | 11 (48%) | 0 (0%) | 2 (22%) |  |  | 6 (46%) | 0 (0%) | 0 (0%) |  |  |
| above | 20 (51%) | 14 (61%) |  |  | 12 (33%) | 10 (83%) | 12 (86%) |  |  | 7 (30%) | 6 (86%) | 7 (78%) |  |  | 5 (38%) | 4 (80%) | 5 (100%) |  |  |
| pp Weight Retained | 3.1 (-0.2, 7.6) | 3.2 (2.0, 8.8) | 0.3 | 0.6 | 1.8 (0.3, 5.0) | 4.7 (1.4, 8.1) | 7.6 (3.1, 11.9) | 0.053 | 0.14 | 1.6 (-1.3, 3.8) | 4.4 (1.0, 6.5) | 7.8 (3.1, 12.4) | 0.083 | 0.2 | 2.5 (1.3, 7.4) | 8.0 (2.7, 8.7) | 6.0 (3.2, 9.8) | 0.5 | >0.9 |
| Length of gestation (days) | 282 (274, 284) | 277 (273, 282) | 0.3 | 0.6 | 280 (273, 284) | 276 (272, 284) | 279 (275, 288) | 0.7 | 0.8 | 282 (276, 284) | 273 (270, 283) | 282 (277, 289) | 0.5 | 0.6 | 277 (270, 282) | 276 (275, 284) | 278 (274, 279) | 0.9 | >0.9 |
| Delivery mode = vaginal | 32 (80%) | 18 (78%) | 0.9 | >0.9 | 30 (81%) | 10 (83%) | 10 (71%) | 0.7 | 0.8 | 20 (83%) | 5 (71%) | 7 (78%) | 0.8 | 0.9 | 10 (77%) | 5 (100%) | 3 (60%) | 0.3 | 0.7 |

<sup>†</sup> n; Mean (SD); n (%); Median (IQR)<sup>2</sup> Kruskal-Wallis rank sum test<sup>3</sup> False discovery rate correction for multiple testing

**Extended Data Table 2.** Infant characteristics.

| Characteristic | Parity |  |  |  | Overall GWG |  |  |  | Primiparous GWG |  |  |  | Multiparous GWG |  |  |  |
| --- | --- | --- | --- | --- | --- | --- | --- | --- | --- | --- | --- | --- | --- | --- | --- | --- |
|  | primiparous<sup>1</sup> | multiparous<sup>1</sup> | p<sup>2</sup> | q<sup>3</sup> | within<sup>1</sup> | above<sup>1</sup> | p<sup>2</sup> | q-value<sup>3</sup> | within<sup>1</sup> | above<sup>1</sup> | p<sup>2</sup> | q-value<sup>3</sup> | within<sup>1</sup> | above<sup>1</sup> | p<sup>2</sup> | q-value<sup>3</sup> |
| n | 24 | 13 |  |  | 12 | 20 |  |  | 7 | 12 |  |  | 5 | 8 |  |  |
| Age | 30.4 (4.3) | 32.7 (3.5) | 0.073 | 0.3 | 32.0 (3.6) | 31.1 (4.0) | 0.5 | >0.9 | 30.6 (3.7) | 30.7 (4.2) | 0.9 | >0.9 | 34.0 (2.4) | 31.9 (3.9) | 0.3 | 0.5 |
| pBMI | 24.2 (20.7, 29.1) | 23.5 (21.5, 25.9) | 0.6 | 0.7 | 21.4 (18.6, 23.6) | 25.8 (24.2, 30.5) | 0.002 | **0.011** | 18.7 (18.6, 25.4) | 25.8 (24.2, 31.1) | 0.031 | 0.2 | 21.5 (21.3, 23.3) | 25.0 (23.9, 27.7) | 0.019 | 0.076 |
| GWG (kg) | 15.0 (11.5, 19.0) | 16.0 (14.0, 20.0) | 0.4 | 0.7 | 13.5 (13.0, 15.0) | 19.0 (17.0, 20.2) | <0.001 | **0.002** | 13.0 (12.0, 15.0) | 19.0 (17.8, 20.2) | 0.003 | **0.036** | 14.0 (13.0, 15.0) | 18.8 (16.8, 21.2) | 0.027 | 0.076 |
| Length of gestation (days) | 281 (276, 284) | 280 (277, 284) | 0.8 | 0.8 | 282 (275, 282) | 281 (278, 288) | 0.4 | >0.9 | 282 (282, 283) | 280 (276, 287) | 0.7 | 0.9 | 277.0 (270.0, 280.0) | 283.0 (279.0, 289.0) | 0.078 | 0.14 |
| Delivery mode = vaginal | 22 (92%) | 13 (100%) | 0.3 | 0.6 | 11 (92%) | 19 (95%) | 0.7 | >0.9 | 6 (86%) | 11 (92%) | 0.7 | 0.9 | 5 (100%) | 8 (100%) |  |  |
| Infant sex = female | 14 (58%) | 5 (38%) | 0.3 | 0.6 | 5 (42%) | 12 (60%) | 0.3 | >0.9 | 4 (57%) | 8 (67%) | 0.7 | 0.9 | 1 (20%) | 4 (50%) | 0.3 | 0.5 |
| Birth weight (g) | 3,431 (447) | 3,655 (353) | 0.11 | 0.3 | 3,540 (306) | 3,526 (490) | >0.9 | >0.9 | 3,665 (246) | 3,320 (510) | 0.069 | 0.3 | 3,366 (318) | 3,835 (245) | 0.028 | 0.076 |
| Birth length (cm) | 51.83 (2.62) | 51.38 (2.26) | 0.5 | 0.7 | 52.17 (2.48) | 51.60 (2.58) | 0.6 | >0.9 | 53.29 (2.75) | 51.42 (2.54) | 0.2 | 0.3 | 50.60 (0.55) | 51.88 (2.80) | 0.4 | 0.5 |
| Birth weight:length | 65 (61, 70) | 72 (70, 74) | 0.024 | 0.3 | 68 (64, 71) | 71 (61, 74) | 0.8 | >0.9 | 69.0 (65.5, 71.6) | 61.7 (59.9, 66.7) | 0.10 | 0.3 | 66.6 (64.7, 69.6) | 72.9 (72.2, 74.9) | 0.023 | 0.076 |
| Birth weight percentile | 50 (20, 60) | 61 (50, 80) | 0.073 | 0.3 | 57 (51, 62) | 54 (20, 80) | 0.8 | >0.9 | 59 (52, 62) | 26 (12, 60) | 0.11 | 0.3 | 46 (28, 58) | 72 (57, 82) | 0.062 | 0.14 |
| Birth length percentile | 46 (18, 74) | 34 (23, 46) | 0.4 | 0.7 | 41 (30, 78) | 46 (18, 62) | 0.5 | >0.9 | 72 (42, 89) | 49 (18, 68) | 0.3 | 0.5 | 31 (24, 38) | 38 (20, 52) | 0.6 | 0.6 |
| Infant weight at 6 months (g) | 7,730 (6,960, 8,250) | 7,738 (7,632, 8,154) | 0.5 | 0.7 | 7,800 (7,305, 8,280) | 7,815 (7,611, 8,200) | >0.9 | >0.9 | 7,730 (6,960, 8,450) | 7,940 (7,634, 8,200) | >0.9 | >0.9 | 7,959 (7,752, 8,154) | 7,662 (7,598, 8,342) | 0.6 | 0.6 |

<sup>1</sup> n; Mean (SD); Median (IQR); n (%) <sup>2</sup> Kruskal-Wallis rank sum test <sup>3</sup> False discovery rate correction for multiple testing

## Notes

### Competing Interest Statement

The authors have declared no competing interest.

### Summary of Updates

This version reflect revisions prior to submission to another journal.

